# Fluralaner treatment of chickens kills the southern house mosquito, *Culex quinquefasciatus*

**DOI:** 10.1101/2024.08.30.610151

**Authors:** Koyle Knape, Yuexun Tian, Cassandra Durden, Dayvion R. Adams, Macie Garza, John B. Carey, Sarah A. Hamer, Gabriel L. Hamer

**Affiliations:** Department of Poultry Science, Texas A&M University; Department of Entomology, Texas A&M University; Department of Veterinary Integrative Biosciences, Texas A&M University; Department of Entomology and Plant Pathology, North Carolina State University, Raleigh, NC, USA

**Keywords:** Xenointoxication, chicken, West Nile virus, systemic insecticide

## Abstract

The control of zoonotic and vector-borne pathogens is challenging due to the limited availability of intervention tools. West Nile virus (WNV) is an example of globally distributed zoonotic arbovirus that circulates between *Culex* species mosquitoes and avian hosts, with spillover transmission to humans, resulting in disease cases. Interventions delivering systemic insecticides to vertebrate hosts used by vector species, known as xenointoxication, are potential tools for managing vector populations by creating toxic bloodmeals. In this study, we evaluated the impact of three systemic insecticides on the mortality of *Cx. quinquefasciatus*: fenbendazole (Safe-Guard® Aquasol), ivermectin (Ivomec® Pour-On), and fluralaner (Bravecto®). We found no significant difference in the feeding rates of mosquitoes that fed on treated chickens compared to those fed on untreated chickens, suggesting that the treatment did not repel mosquitoes. The mortality of *Cx. quinquefasciatus* mosquitoes feeding on fluralaner-treated chickens was significantly higher (p < 0.01) than those fed on control chickens at 3 and 7 days post-treatment, but this effect was not observed in mosquitoes fed on chickens treated with fenbendazole or ivermectin. No differences in mortality were observed among the groups at 14, 26, or 56 days post-treatment. These data support fluralaner as a xenointoxication tool to control *Cx. quinquefasciatus* populations and decrease the risk of human exposure to their associated pathogens.

## 1. Introduction

Zoonotic and vector-borne pathogens are emerging globally, and current control tools are insufficient. Given the many host animals involved, it is particularly difficult to manage the amplification of zoonotic agents among domestic or wild animals with spillover to humans. A pathosystem in this category is West Nile virus (WNV), one of the most widely distributed zoonotic, arthropod-borne viruses in the world. It is primarily transmitted by *Culex* spp. mosquitoes after blood feeding on viremic avian amplification hosts (McLean et al. 2001). In the United States, WNV vector varies in different regions with *Cx. tarsalis* leading transmission in western states, *Cx. quinquefasciatus* in southern states, and *Cx. pipiens* in northern states (Ciota 2017). Since the introduction of WNV to New York City in 1999, WNV has become widespread and the most common human mosquito-borne pathogen in the United States (Soto et al. 2022), with over 7 million human cases (Ronca et al. 2021). In the U.S., approximately 1 out of 150 WNV-infected people develop severe neurological illness and functional sequelae. Among those with severe illness, the case-fatality ratio is approximately 10% (CDC 2024) generating an estimated $56 million U.S. dollars in annual medical expenses (Ronca et al. 2021).

Due to a lack of commercially available vaccines, controlling WNV is primarily achieved by population suppression of *Culex* spp. mosquitoes through larval source reduction, larvicides, adulticides, and public education to prevent vector bites (Nasci et al. 2013). In addition to its high cost, using insecticide to control mosquitoes raises concerns about impacts on human health (Ross et al. 2013), effects on non-target invertebrates (Rasmussen et al. 2013), and insecticide resistance development. This highlights the need to develop novel approaches for insecticides application to manage WNV. Host-targeted (systemic) insecticides, such as ectoparasiticides and endectocides (e.g., lethal on ecto-and endoparasites) drugs, offer strategies for controlling human malaria (Foy et al. 2011), Chagas disease (Dias et al. 2005), and more recently, WNV (Nguyen et al. 2019). Ivermectin, a broad-spectrum endectocide, provided as treated seeds via artificial feeders to free-ranging passerine birds resulted in lethal bloodmeals that kill WNV vectors (Nguyen et al. 2019, Holcomb et al. 2022). However, ivermectin reaches maximum concentration immediately after treatment and may be undetectable within 24 hours (Arisova 2020). Similarly, Nguyen et al. (2019) found levels of ivermectin in chicken serum dropped quickly, with observed mosquito mortality only until 3 days post-treatment. The need to repeatedly treat hosts with systemic insecticides could be accomplished using treated feed yet limits the feasibility of scale-up for population control of vectors. Accordingly, the use of alternative active ingredients deliver toxic bloodmeals to mosquito vectors for longer durations is desirable for host-targeted vector control interventions. One alternative active ingredient is fluralaner, which was recently shown to produce toxic bloodmeals to *Aedes aegypti* mosquitoes for up to 15 weeks post-treatment in dogs (Evans et al. 2023) and to triatomines for up to 14 days post-treatment in chickens (Durden et al. 2023). It has been proposed to be used to control vector-borne human diseases (Miglianico et al. 2018).

*Culex* mosquito vectors of WNV in the United States regularly utilize avian hosts for bloodmeals (Hannon et al. 2019). While ongoing studies are using host-targeted insecticide treatment of wild granivorous birds (Holcomb et al. 2023), the primary vector of WNV in the southern half of the US, Mexico, and Central America is *Culex quinquefasciatus*, which was documented to feed on chickens 67% of the time in South Texas (Olson et al. 2020), 45.3% in Guatemala (Kent et al. 2010), and 6.1% in Reynosa, Mexico (Estrada-Franco et al. 2020). Chickens are ubiquitous in the peridomestic environment in some regions, which offers a convenient target for creating toxic bloodmeals by systemic insecticides and are easier to treat than wild passerines. Additionally, sentinel chicken flocks are routinely used as a surveillance tool to detect WNV circulation by detecting seroconversion (Chaskopoulou et al. 2013). Therefore, xenointoxication with chickens is a potential tool for the wide-scale population suppression of *Culex* mosquitoes to reduce the risk of human and animal exposure to WNV.

In this study, we evaluate the off-label, systemic treatment of chickens with three commercially available products to evaluate their insecticidal efficacy on *Cx. quinquefasciatus* when fed directly treated chickens. We tested three active ingredients (fenbendazole, ivermectin, and fluralaner). To our knowledge, this study is the first to document the utility of fluralaner systemic treatment of chickens for the control of mosquitoes.

## 2. Materials and Methods

### 2.1 Mosquito colony

*Culex quinquefasciatus* Sebring strain (SEB) was used in this study (Sbrana et al. 2005). Mosquitoes were maintained on a natural day and night light cycle (10 h light, 14 h dark) with a constant 50% humidity at 27 °C and provided a 10% sucrose solution. Maintenance feeding of the colony was comprised of whole chicken blood treated with heparin (Sagent Pharmaceuticals, Schaumburg, IL) using Hemotek membrane feeders (Hemotek, Ltd., Blackburn, UK).

### 2.2 Chicken

A flock of laying chickens (*Gallus gallus domesticus*) was obtained from a commercial hatchery (Lohmann LSL-Lite, Cuxhaven, Germany). These chickens were enrolled in this study at 28 weeks of age and were confirmed as healthy based on daily clinical health evaluations, egg-laying records, and body weight records. Chickens were housed in an environmentally controlled layer house with a light/dark cycle of 16h/8h, which is suitable for egg production. Within the laying house, the chickens were housed with two per cage with a nipple waterer that serves two cages (four chickens) and fed on a standard commercial layer diet before experiments.

### 2.3 Chicken Treatment

Chicken treatment was previously described in Durden et al. (2023). Briefly, the individual chicken was treated with one of the three systemic insecticides: ivermectin (Ivomec®, Boehringer Ingelheim, Ingelheim am Rhein, Germany), fenbendazole (Safe-Guard® AquaSol, Merck Animal Health USA, Rahway, NJ), and fluralaner (Bravecto®, Merck, Rahway, NJ, USA), or regular food/water as a control. The dose of each insecticide was calculated based on the average hen weight (1.34 kg). Ivermectin and fenbendazole were delivered to chickens in a liquid formulation using a gravity flow nipple watering system with the chemical mixed into their water and dosed at 0.4 mg/kg of body weight for ivermectin (Holcomb et al. 2022) and 1 mg/kg for fenbendazole as indicated on the product label. Fluralaner was delivered as a small oral chew in a dose of 0.5 mg/kg (Thomas et al. 2017) before the daily food was provided to ensure a full consumption of the oral chew. Ivermectin and fenbendazole treatments were conducted daily for five consecutive days (Nguyen et al. 2019), while fluralaner was given to the chickens twice, seven days apart (Thomas et al. 2017).

### 2.5 Mosquito Feeding

Experiments were carried out using 7-12 days old adult *Cx. quinquefasciatus* females, which were starved for 24 hours before the experiment. During each trial, 42-99 mosquitoes were used to feed on a treated or control chicken at days 3, 7, 14, 28, and 56 days post-treatment (DPT). Each chicken was used only once to avoid confounding effects of acquired immunity to mosquito salivary antigens. Mosquitoes were briefly knocked down by placing their BugDorm (MegaView Science Co., Ltd., Taichung, Taiwan) into a -20°C freezer for 5-10 minutes to sort females. The females were placed into a plastic container with a square opening (10×10 cm) on the side to which a mesh sleeve was glued on. The chicken’s feet were inserted into the mosquito container through the mesh sleeve for mosquitoes to feed on for 45 min, while the chicken was restrained using a stretchable self-adherent wrap (Healqu, Jersey City, New Jersey) on a padded stainless steel oven grate. The same chicken had three *T. gerstaeckeri* feeding on the breast at the same time as described in Durden et al. (2023). After the 45-minute feeding period, the mosquitoes were knocked down immediately by putting the container into a cooler (61L x 38H x 61W cm) half-filled with ice for 15 minutes. The immobilized mosquitoes were sorted with unfed mosquitoes removed and blood-fed mosquitoes were transferred into a new container (5,342 cm^3^) with a 10% sucrose solution. The container was then placed into an incubator at 27 °C and 50% humidity to monitor the survivorship of the blood-fed mosquitoes, which was checked daily for 10 days. The trial was repeated three times resulting in a total of 60 chickens ([3 treatment + 1 control] * 5 timepoints * 3 replicates) used in this study.

### 2.7 Statistics

All statistical analyses were conducted using R studio software (Version 4.2.2R Foundation for Statistical Computing, Vienna, Austria). Feeding success was calculated by dividing the number of blood fed mosquitoes by the total number of mosquitoes in the container. The effects of treatment and DPT were analyzed using Analysis of Variance (ANOVA) followed by Tukey’s post-hoc test (R package: stats. (Team 2020)). Mosquito survivorship was analyzed using the Kaplan-Meier survival curve (R package: survival (Therneau 2020)) followed by a paired log-rank test (R package: survminer (Kassambara et al. 2021)) to compare the survival curves with different treatments.

## 3. Results

### 3.1 Feeding Success

There was no significant difference detected among the treatments within each DPT or among the DPT within each treatment (Figure 1). However, a significant difference was observed under different DPT (14 vs 56, *p*-value = 0.013), while no significant difference was detected between the treatments and the interaction of DPT and treatment.

**Figure 1.**
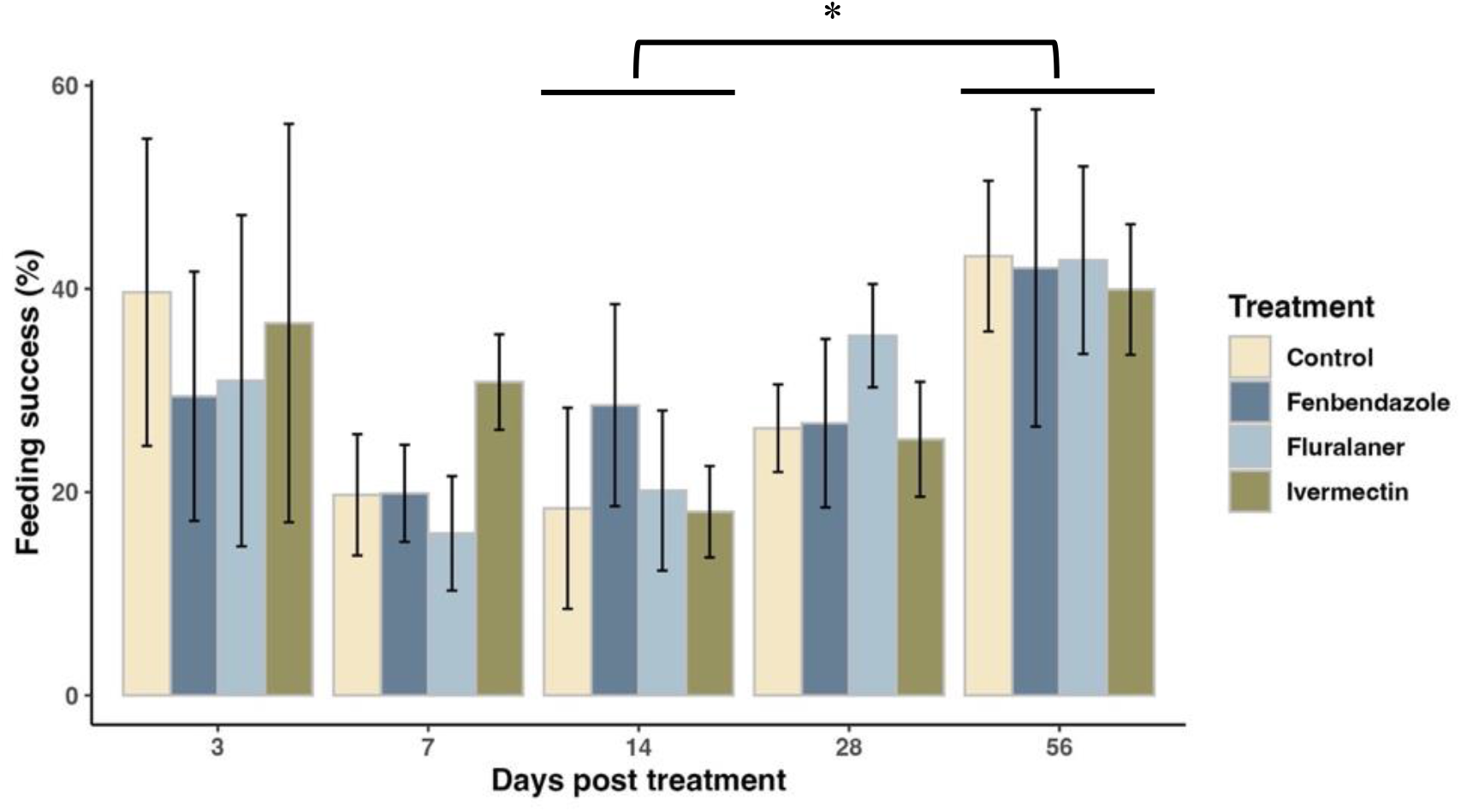
The mean (± SE) of *Culex quinquefasciatus* feeding success on chickens with different treatments at different time points post-treatment.

### 3.2 Survivorship

Significant differences in survivorship were observed only at 3, 7, and 56 DPT, but not at 14 and 28 DPT (Figure 2). At 3 DPT, the mosquito survivorship with fluralaner was significantly lower compared to the control (*p*-value < 0.001), while mosquitoes fed on ivermectin (*p*-value = 0.0012) and fenbendazole-treated chickens (*p*-value = 0.0002) had higher survivorship than the control. The same result was observed for fluralaner at 7 DPT (*p*-value = 0.0042), but ivermectin (*p*-value = 0.4767) and fenbendazole-treated chickens (*p*-value = 0.4770) were not significantly different at this time point. At 56 DPT, mosquitoes with fenbendazole had a significantly higher survivorship than mosquitoes in the control group (*p*-value = 0.0137), but no significant difference were observed for ivermectin (*p*-value = 0.8666) and fluralaner (*p*-value = 0.8666).

**Figure 2.**
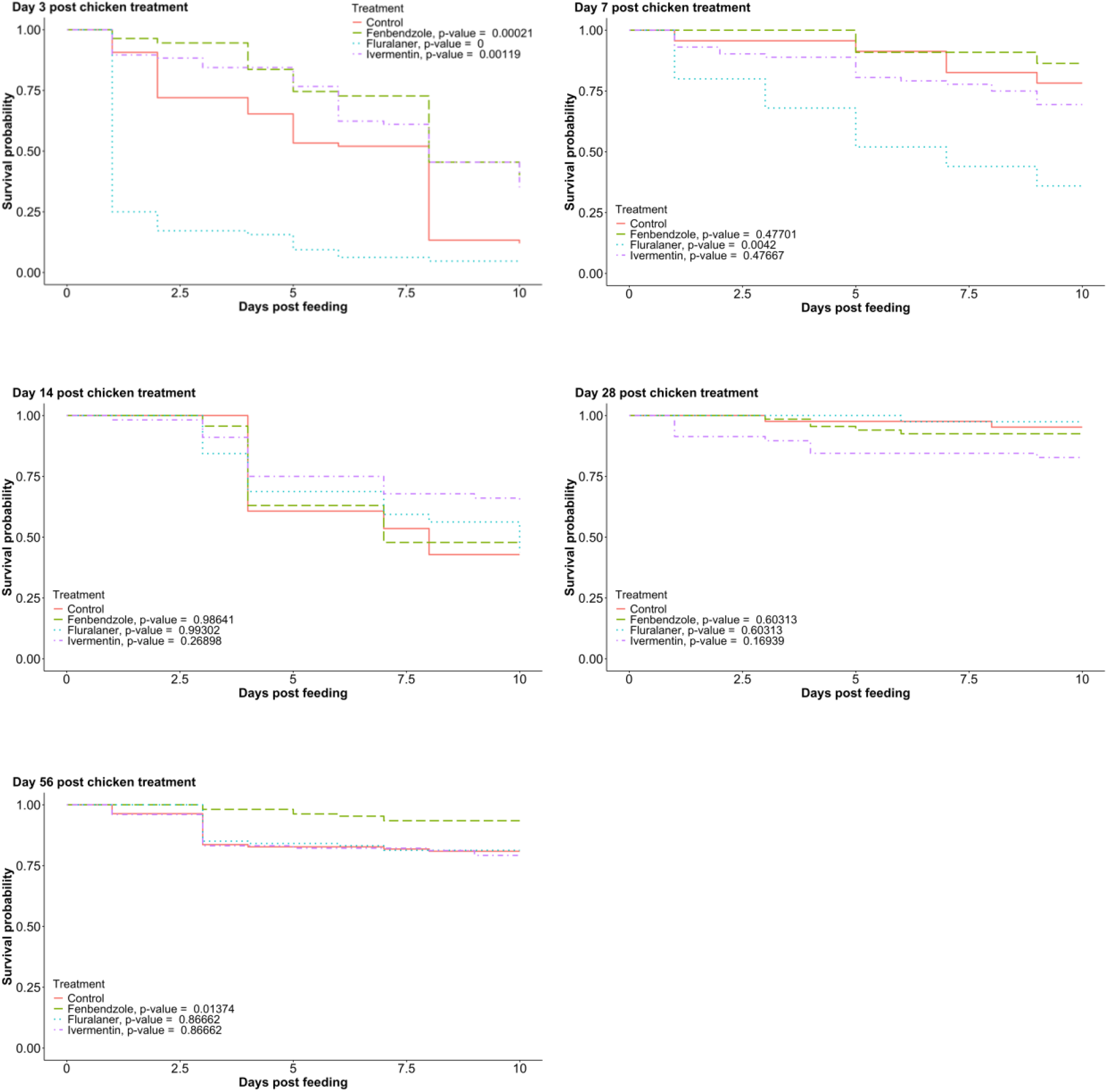
Kaplan-Meier survival curve of mosquitoes fed on chickens as control or chickens treated with fenbendazole, ivermectin, or fluralaner at 3, 7, 14, 28, and 56 days post-treatment.

## 4. Discussion

In this study, we evaluated the mortality of *Cx. quinquefasciatus* after feeding on chickens that were treated with three systematic insecticides including fluralaner, ivermectin, and fenbendazole, for up to 56 days post-treatment. Only fluralaner-treated chickens resulted in significant mortality in mosquitoes compared to the control group at least 7 DPT, but not 14 or longer DPT. These results were consistent with Durden et al. (2023) showing that fluralaner, but not ivermectin or fenbendazole, was detectable in chicken plasma at 7 and 14 DPT.

Evans et al. (2023) and Duncan et al. (2023) documented a longer effective period, up to 15 and 12 weeks respectively, against *Aedes aegypti* from fluralaner (Bravecto)-treated dogs with fluralaner detected from dog plasma at all timepoints in Evans et al. (2023). The differences between the effective periods may be due to the doses administered by chickens and dogs. Bravecto®, as a commercial product for dogs to treat fleas and ticks (*Ixodes scapularis, Dermacentor variabilis, Rhipicephalus sanguineus*, and *Haemaphysalis longicornis*), contains the optimal doses for dog treatment, but not for chickens. Previous studies have identified the lethal dose concentrations (LD_50_) of fluralaner to *Ae. aegypti* and *Cx. quinquefasciatus* to be 24.04 and 49.82 ng/mL respectively, based on artificial inoculations of heparinized chicken blood (Shah et al. 2024). In our study, this LC_50_ for *Cx. quinquefasciatus* was only achieved in chicken plasma for up to 7 DPT (Durden et al. 2023), which supports our observed mosquito survival results. Fluralaner has been previously shown to be safe in chickens up to 5 times the dose used in our study (Prohaczik et al. 2017), so exploring the optimal dose of fluralaner for chicken treatment to achieve a longer effective period against mosquitoes is necessary. In Europe, fluralaner has been commercialized as Exzolt™ (Merck Animal Health USA, Rahway, NJ) to treat poultry mites in a liquid formulation with 0-day egg withdrawal and 14-day meat withdrawal according to the product manual. However, it has not yet been approved in the US and there are no other commercial fluralaner products available for poultry.

Ivermectin has been used as an endectocide to treat nematodes and ectoparasites on domestic animals (Foy et al. 2011) with high lethal and sublethal effects on mosquitoes such as *Anopheles* spp. (Foley et al. 2000, Derua et al. 2016), *Aedes* spp. (Deus et al. 2012), and *Culex* spp. (Deus et al. 2012, Nguyen et al. 2019). However, its short half-life limits the effects of arthropod vector control. Nguyen et al. (2019) reported that ivermectin concentration in chicken serum rapidly decreased after consumption and was only detectable up to two days, which was positively correlated to *Cx. tarsalis* mortality that was fed on those treated chickens. Our results are consistent with the observations of Nguyen et al. (2019) as the earliest mosquito feeding was conducted on 3 DPT and survivorship of mosquitoes fed on ivermectin-fed chickens was not lower than those fed on control birds. This result indicates the need for repeated chicken treatments to achieve long-term management of *Cx. quinquefasciatus*, which increases control costs.

Fenbendazole is an antiparasitic used as the active ingredient in the commercial product Safe-Guard® AquaSol (Merck Animal Health USA, Rahway, NJ) which is labeled for parasite treatment for chickens. Derouen et al. (2009) evaluated the effects of Safe-Guard® (fenbendazole, cattle formulation) on gastrointestinal nematodes in calves, which were treated at a dose of 5 mg/kg. The treatment of fenbendazole significantly reduced the nematode eggs in feces with a reduction rate of 96-100%, suggesting that fenbendazole was effective in controlling nematode infestation in calves (Derouen et al. 2009). However, few studies have evaluated the effects of fenbendazole on arthropod feeding. In this study, fenbendazole had no significant effect on mosquito mortality, indicating either a lack of insecticidal effects or that the minimum dose was not reached in chicken blood to kill mosquitoes, which is consistent in *Triatoma gerstaeckeri* (Durden et al. 2023). Further research is needed across dose ranges to confirm the effects of fenbendazole on mosquitoes, triatomines, and other ectoparasites.

Overall, less than 50% of the mosquitoes fed on treated or control chickens. This low feeding percentage is consistent with prior studies demonstrating low feeding rates in laboratory environments by *Culex* sp., while other colonized mosquito species such as *Aedes aegypti*, are more willing to take bloodmeals (Lyski et al. 2011, Meuti et al. 2023). Our low feeding rates could also be attributed to other factors. For example, although the chickens were restrained, their movements during feeding may have still prevented bloodmeal acquisition. Mosquitoes may also be damaged or stressed while transferring in vehicles from the insectary to the experimental feeding room 2km away. Despite the low feeding rate, no significant differences were observed within each time point or each treatment, suggesting no repellent effects of the products evaluated in this study on mosquitoes.

The results reported in our study and previous studies (Alcantara et al. 2023, Duncan et al. 2023, Durden et al. 2023, Evans et al. 2023) suggest fluralaner is a promising candidate as a host-targeted insecticide due to high efficacy against blood-feeding arthropods, lack of repellency, and non-toxicity in a wide range of domestic animals. This approach of xenointoxication could allow for area-wide treatment of chickens in peridomestic environments which could result in population suppression of medically relevant bird-biting mosquitoes such as *Cx. quinquefasciatus*.

*Culex quinquefasciatus* is nearly globally distributed and considered to be an important vector of multiple pathogens including West Nile virus (Ciota 2017), St. Louis encephalitis virus (Diaz et al. 2013), and filarial nematodes such as *Wuchereria bancrofti* (Calheiros et al. 1998). Xenointoxication is an alternative mosquito control tool that could achieve population suppression of a variety of blood feeding arthropods of public health importance and future studies should evaluate the impact of xenointoxication on disease transmission in nature.

## Acknowledgments

We appreciate the facility support from Texas A&M University Veterinary Medicine & Biomedical Science Veterinary Medical Park.

## Ethics approval and consent to participate

Protocols for the use of chickens was reviewed and approved by the Texas A&M University Institutional Animal Care and Use Committee (Animal Use Protocol IACUC 2021-0109) on 05/11/2021.

## Author contribution statement

Conceptualization: GH, DA, SH; Methodology: GH, CD, KK, JC; Data curation: CD, KK, MG; Formal Analysis: YT; Writing-Original Draft Preparation: KK, YT; Writing – Review & Editing: all authors.

